# A closed *Candidatus* Odinarchaeum genome exposes Asgard archaeal viruses

**DOI:** 10.1101/2021.09.01.458545

**Authors:** Daniel Tamarit, Eva F. Caceres, Mart Krupovic, Reindert Nijland, Laura Eme, Nicholas P. Robinson, Thijs J. G. Ettema

## Abstract

Asgard archaea have recently been identified as the closest archaeal relatives of eukaryotes. Their ecology remains enigmatic, and their virome, completely unknown. Here, we describe the closed genome of *Ca*. Odinarchaeum yellowstonii LCB_4, and, from this, obtain novel CRISPR arrays with spacer targets to several viral contigs. We find related viruses in sequence data from thermophilic environments and in the genomes of diverse prokaryotes, including other Asgard archaea. These novel viruses open research avenues into the ecology and evolution of Asgard archaea.

---

Asgard archaea are a diverse group of microorganisms that comprise the closest relatives of eukaryotes^1–7^. Their genomes were first explored over six years ago^8^, and much of their physiology and cell biology remains to be studied. While over two hundred draft genomes are available for this group, the majority is represented by highly fragmented and incomplete metagenome assembled genomes (MAGs), which has precluded obtaining insights into their mobile genetic elements (mobilome). Given the central role of Asgard archaea in eukaryogenesis models, access to their complete genomes and information about their interactions with viruses are highly relevant.

To obtain a complete Asgard archaeal genome, we reassembled the Odinarchaeota LCB_4 genome, currently a 96% complete assembly of 1.46 Mbp distributed in 9 contigs^1^. A promising reassembly yielded a 1.41 Mbp contig, a 13 kbp contig containing CRISPR-associated (*cas*) genes, and multiple short contigs harboring mobile element or repeat signatures (see Methods for details). After contig boundary inspection, we postulated that the first two contigs represented the entire Odinarchaeota LCB_4 chromosome as these were flanked by similar CRISPR arrays that extended for several kilobasepairs (Suppl. Fig S1). We successfully amplified these gaps using long-range PCR, sequenced the resulting amplicons with Nanopore sequencing, and performed a hybrid assembly, finally generating a single 1.418 Mbp circular contig. Given the high quality of this genome, we suggest recognizing this strain as *Candidatus* Odinarchaeum yellowstonii LCB_4 (hereafter ‘LCB_4’), in reference to Yellowstone National Park, location of the hot spring where it was sampled.

The LCB_4 genome contains a unique disposition of CRISPR-Cas genes (Fig. 1), including neighboring Type I-A and Type III-D *cas* gene clusters, separated by a 6.1 kb-long Type I-A CRISPR array, and further followed by another 2.7 kb-long Type I-A CRISPR array, with a total of 144 CRISPR spacers across both arrays. Nine of these spacers targeted, with 100% identity and query coverage, four putative mobile element contigs obtained in the same assembly that were not part of the closed genome (Fig. 1). Two of these contigs contained genes encoding common mobile element proteins, such as restriction endonucleases and integrases, but did not contain any viral signature genes (Suppl. Table S1). A third contig represented a complete, circular viral genome (Suppl. Fig. 1E) encoding transcriptional regulators, an endonuclease, and a double jelly-roll major capsid protein (DJR-MCP), typical of tailless icosahedral viruses (Fig. 1; Suppl. Fig. S2A; Suppl. Table S1). This specific protein was previously found in a study of the DJR-MCP family, and was tentatively classified as belonging to an “Odin group” since it was found in the same metagenome as *Ca*. Odinarchaeum LCB_4^9^. The present complete recovery of LCB_4 CRISPR arrays allowed us to confirm that this circular contig indeed represents a virus associated with *Ca*. Odinarchaeum, for which we suggest the name Huginnvirus, in reference to Odin’s raven, Huginn (“thought”).

**Figure 1.**
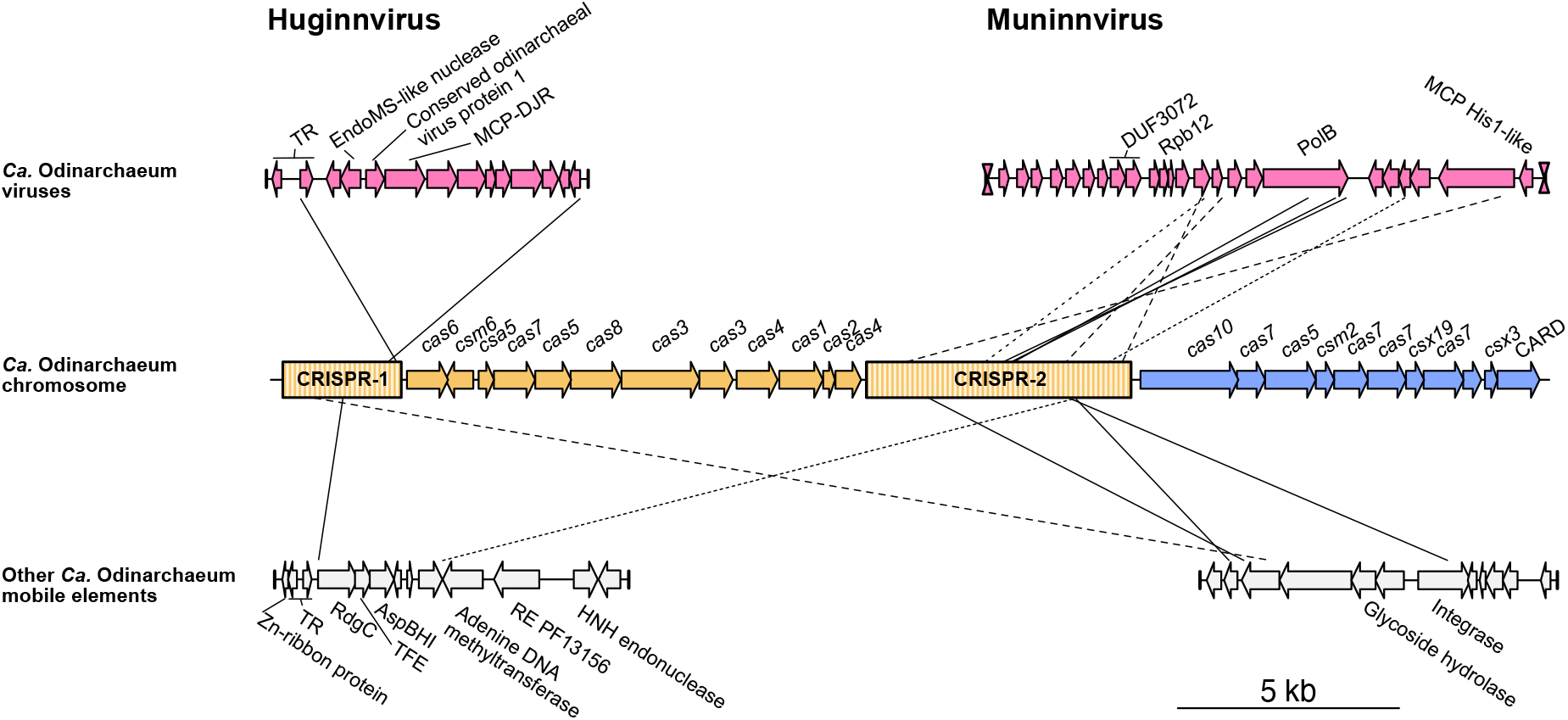
*Ca*. Odinarchaeum LCB_4 CRISPR-Cas system and mobile elements. CRISPR-Cas systems in the *Ca*. Odinarchaeum LCB_4 chromosome (center) were colored according to their type classification (orange: I-A; blue: III-D). Full contigs representing mobile elements are shown at the corners, with vertical lines representing contig boundaries. Viral terminal inverted repeats are represented by hourglass symbols. Connecting lines represent full-coverage spacer (35-42 nt) BlastN hits against mobile element targets (0 mismatches: full; 1 mismatch: dashed; 2 mismatches: dotted). TR=Transcriptional regulator; MCP=Major Capsid Protein; DJR=Double Jelly-Roll; TFE=Transcription Initiation Factor E; RE=Restriction enzyme; PolB=Family B DNA polymerase.

Furthermore, 3 spacers yielded 100% identity (and a further 5 spacers if 1-2 mismatches were permitted) and 100% query coverage matches against a 12.7 kb-long contig recovered by the new *Ca*. Odinarchaeum LCB_4 assembly (Fig. 1). All 3 hits targeted an ORF encoding a primed family B DNA Polymerase (PolB), a gene frequently observed in archaeal viruses. Further inspection of this contig revealed genes encoding a zinc-ribbon protein and a His1-family MCP (Suppl. Fig S2B; Suppl. Table S1), conserved in spindle-shaped viruses^10^. This contig was flanked by ca. 80 nucleotides-long terminal inverted repeats, a typical signature of viruses with linear dsDNA genomes replicated by protein-primed PolBs^11^. Thus, this contig represents a complete Asgard archaeal viral genome for which we suggest the name Muninnvirus, in relation to Odin’s raven, Muninn (“memory”).

We further queried the PolB sequence from the Muninnvirus genome through phylogenetic analysis, finding that it is closely related to a homolog in a Sulfolobus ellipsoid virus 1 (SEV1)^12^ (Fig. 2A), recently isolated from a Costa Rican hot spring. No other genes were found to be shared between Muninnvirus and SEV1, indicative of recent horizontal transfer of *polB* in at least one of these viruses. Interestingly, other close homologs included multiple sequences that were likewise obtained from hot springs or hydrothermal vents (Fig. 2A). Two of these hits were part of an Asgard archaeal MAG (QZMA23B3), and a third one belonged to a MAG (SpSt-845) originally classified as Bathyarchaeota. A phylogenomic analysis that indicated that the QZMA23B3 belonged to the recently described Asgard archaeal class Jordarchaeia^7^, and that SpSt-845 in fact belonged to the Nitrososphaeria (Suppl. Fig. S3). Two additional PolB sequences from the same Nitrososphaerial MAG were found to be highly similar (>80% identity) to HGY28086.1 (and were thus not included in the phylogenetic analysis). The five PolB homologs were encoded in contigs containing SIRV2-famiily MCP genes (Fig. 2B; Suppl. Fig. S2C; Suppl. Table S1), exclusive to archaeal filamentous viruses with linear dsDNA genomes and classified into the realm Adnaviria^13^. Both the Jordarchaeia and the Nitrososphaeria contigs displayed high conservation in synteny and protein sequences, indicating high contig completeness and recent diversification (Fig. 2B). Notably, none of the known archaeal viruses with SIRV2-family MCP encodes its own PolB, suggesting that the group identified herein represents a new archaeal virus family. However, while we detected CRISPR arrays in the MAGs where these viral contigs were identified, we could not find accurate spacer matches (query coverage > 90%, identity > 90%) to these viral sequences and therefore the identity of the hosts of these thermophilic viruses remains unclear.

**Figure 2.**
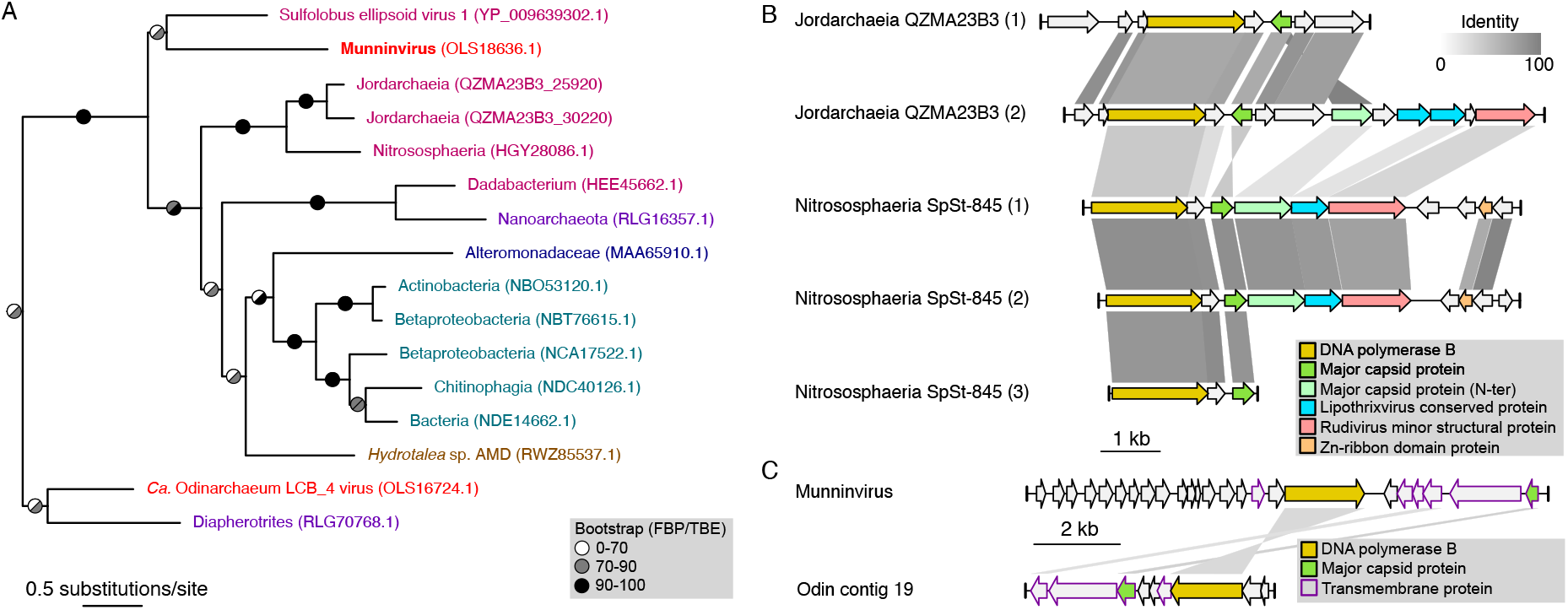
Discovery of additional Asgard archaeal mobile elements. (A) Phylogeny of DNA polymerase B obtained with IQTree2 under the Q.pfam+C60+R4+F+PMSF model. Color: *Ca*. Odinarchaeum LCB_4 MAG (red); sequences obtained from hot springs (pink); hydrothermal vents (purple); marine water (dark blue); Chatahoochee river (USA) (light blue); mine drainage (brown). Branch support value are Felsenstein Bootstrap Proportions (left) and Transfer Bootstrap Expectation (right). The tree presented here is a clade of the full tree shown in Suppl. Fig. S4. (B and C) Comparison between the viral contigs (B) of Jordarchaeia QZMA23B3 and Nitrososphaeria SpSt-845, and (C) of Muninnvirus and another viral contig in the *Ca*. Odinarchaeum LCB_4 assembly. Similarity lines represent BlastP hits with E-value lower than 1e-5, and percent identity as shown in the upper-right legend.

The PolB phylogeny further suggests that a clade of viral sequences found in MAGs from mesophiles evolved from a likely thermophile-infecting ancestor. While none of the mentioned mobile elements share other proteins in common with Muninnvirus, a more distant relative of the Muninnvirus PolB sequence was found in a contig from the same assembly. Like Muninnvirus, this sequence encoded a His1-like MCP, and a gene encoding a transmembrane protein of unknown function (Fig. 2C). The latter two genes surrounded another gene encoding a relatively long protein (580 and 553 amino acid residues) with multiple transmembrane helices and complex predicted structures (Suppl. Fig. S2D), with no detectable similarity, but possibly related functions.

The CRISPR-Cas system of *Ca*. Odinarchaeum yellowstonii LCB_4 is likely its primary antiviral defense system. We could find no homologs for DISARM^14^ or other recently discovered antiviral systems^15,16^ in its genome. The retention of numerous CRISPR spacers against these mobile elements is significant and indicates coevolutionary dynamics with viruses from multiple families.

Our findings highlight the benefits of improving the quality of Asgard archaeal genomes. The discovery of viruses of thermophilic Asgard archaea opens the door to the study of the Asgard archaeal mobilome, promising exciting new insights into the ecology, physiology and evolution of the closest archaeal relatives of eukaryotes.

## ACKNOWLEDGEMENTS

We thank Laura Wenzel for discussions on hybrid assemblies, and Raymond Staals and John van der Oost for helpful comments on CRISPR-Cas systems. This research was funded by the Swedish Research Council (International Postdoc grant 2018-00669 to DT), the European Research Council (ERC consolidator grant 817834 to TJGE), the Swedish Foundation for Strategic Research (SSF-FFL5 to TJGE) and a Wellcome Trust collaborative award (203276/Z/16/Z to TJGE). NR was supported by start-up funds from BLS, Lancaster University and a Leverhulme Research Project Grant (RPG-2019-297). MK was supported by the Agence Nationale de la Recherche (ANR-20-CE20-0009-02) and Ville de Paris (Emergence(s) project MEMREMA). LE received funding from the European Research Council (ERC Starting Grant 803151).

## DATA AVAILABILITY

Raw Nanopore amplicon reads and complete *Ca*. Odinarchaeum LCB_4 assembly are available at NCBI under Bioproject PRJNAXXXXXXX. Additional data and supporting alignments and trees can be found in Zenodo, under the link: XXXXXXXXX.

## METHODS

### Odinarchaeon LCB_4 genome reassembly

To reassemble the Odinarchaeon LCB_4 genome (Suppl. Fig. S1A), its corresponding Illumina reads^17^ (BioSample SAMN04386028) were mapped against Asgard MAGs^4^ using Minimap2^18^ v2.2.17. Mapped reads were extracted and assembled with Unicycler^19^ v0.4.4. This assembly obtained a 1.406 Mb contig, which was not predicted as circular despite both of its contig boundaries ending in CRISPR arrays (Suppl. Fig. S1B). Additional short (< 13 kb) contigs were not considered part of the main chromosome if they represented mobile elements (with signatures such as differing coverage, circularity, CRISPR spacer hits and/or presence of typical mobile element genes), rRNA genes from different organisms, or CRISPR arrays. The latter two were expected due to the conservation of sequences such as rRNA gene sequences and CRISPR repeats. After removing these contigs, only one additional contig of 10.6 kb containing type I-A *cas* genes remained. Given that the 1.406 Mb contig ended in type I-A CRISPR arrays, we hypothesized that these two contigs could represent the entire circular chromosome of *Ca*. Odinarchaeum LCB_4. In parallel, we assembled the Illumina reads with MEGAHIT^20^ v.1.1.3 with options “--k-min 57 --k-max 147 --k-step 12”. While highly fractionated, this assembly found an alternative solution for the sequences involved in the contig borders of the previous assembly (Suppl. Fig. S1B). In this case, the type-I-A *cas* genes were surrounded by two separate CRISPR arrays. Moreover, four consecutive spacers in the innermost side of one of the CRISPR arrays in this assembly were identical to the outermost spacers of the CRISPR array present at the border of the 1.406 Mb contig in the Unicycler assembly (Suppl. Fig. S1B). These results suggested a specific disposition for the two aforementioned contigs.

### Long-range PCR and Nanopore sequencing

Four regions were selected for long-range Polymerase Chain Reaction (lrPCR): two contig gaps, corresponding to CRISPR arrays, and two control regions spanning ca. 5 kb of the rRNA operon and ca. 10 kb of a ribosomal protein gene cluster (Suppl. Table S2). Primers were designed using the Sigma-Aldrich OligoEvaluator™ (http://www.oligoevaluator.com/OligoCalcServlet) and synthesized by Integrated DNA Technologies, Inc. MDA-amplified environmental DNA isolated from the Lower Culex Basin at Yellowstone National Park (USA)^17^ was then amplified with Herculase polymerase (Agilent). Amplification of control and gap regions was then performed following the parameters shown in Suppl. Table S2. Products were separated on a 0.8% agarose gel in 1XTBE stained with SYBR-Gold, and purified using a Qiagen Spin purification kit following the manufacturers instructions. Purified PCR fragments were pooled and used to construct a library with the ligation kit SQK-LSK109. Sequencing was performed on an Oxford Nanopore MinION Mk1C sequencer using an R9.4.1 flow cell. Raw sequence data was basecalled using Guppy v4.2.2 High Accuracy. Reads were separated in two bins at 3-9 kb (subsampled to 30X) and 9-12 kb, and processed to obtain consensus sequences using Decona (https://github.com/Saskia-Oosterbroek/decona) v0.1.2 (-c 0.85 -w 6 -i -n 25 -M -r). Both control regions, comprising the rRNA and ribosomal protein operons, were 100% identical to the corresponding nucleotide sequences of the published assembly.

### Hybrid assembly

Reads were filtered using NanoFilt v.2.6.0 with options “-q 10 -l 1000”. We used these filtered Nanopore reads and the mapped Illumina reads to perform a hybrid assembly with Unicycler 0.4.4, which resolved both the main chromosomal contig and a viral contig (Huginnvirus) as circular (Suppl. Fig S1DE). Read mapping was performed using bowtie2^21^ v.2.3.5.1 for Illumina reads and minimap2^18^ v.2.17.r941 for Nanopore reads. A local cumulative GC skew minimum (Suppl. Fig S1E), together with low R-Y (purine minus pyrimidine), M-K (amino minus keto) and cumulative AT skew values, was selected as a potential replication origin and the circular contig was permutated to set this position as nucleotide +1.

### Annotation

CRISPR arrays were detected and classified using CRISPRdetect^22^ v2.4 online, and *cas* genes were detected and classified through CRISPRcasIdentifier^23^ v1.1.0. Proteins were classified into COG families^24^ based on 5 best local Blastp^25^ hits to the same COG, and domain annotation was performed through InterProScan^26^ v5.48-83.0. Mobile element protein annotation was performed using HHsearch^27^ against Pfam^28^, PDB^29^, SCOPe^30^, CDD^31^ and Uniprot^32^ viral protein sequence databases. Synteny plots were performed with genoPlotR^33^. Structural predictions were performed with RoseTTAFold^34^ through the Robetta portal.

### Phylogenetics

Reference DNA polymerase B sequences were obtained from Kim et al.^35^ and used for Psiblast^36^ v2.10.0+ against the NR database and a group of Asgard archaeal MAGs from Eme et al.^4^. Sequences with over 70% similarity were removed with CD-Hit^37^ v4.7. The remaining sequences were aligned with Mafft-linsi^38^ v7.450 and columns with over 50% gaps were removed using trimAl^39^ v1.4.rev22. Additionally, sequences with over 50% gaps in the trimmed alignment were removed. Maximum-likelihood trees were reconstructed using IQ-TREE^40^ v2.0-rc1 using ModelFinder^41^ with all combinations of the empirical models LG, JTT, WAG and Q.pfam with site-class mixtures (none, C20, C40, C60), rate heterogeneity (none, G4 and R4) and frequency (none, F) parameters. Using the previous tree as a guide, a PMSF^42^ approximation of the selected model was used to reconstruct a tree with 100 non-parametric bootstrap pseudo-replicates, which was then interpreted both as the standard Felsenstein bootstrap proportions and as transfer bootstrap expectation^43^.

To assess the taxonomy of selected MAGs, all archaeal GTDB^44^ representative sequences (as of August 2021) were retrieved and supplemented with Asgard archaeal sequences from the Hermod^45^, Sif^5^, Wukong^6^ and Jord^7^ groups. Together with query sequences, GToTree^46^ v1.5.45 was then used to reconstruct a tree with parameters “-H Archaea -D -G 0.2”.

## SUPPLEMENTARY MATERIAL

**Supplementary Table S1. Annotation of mobile elements.** Yellow cells represent annotation of key viral proteins.

**Supplementary Table S2. Methods: primers and lrPCR cycling parameters**

**Supplementary Figure S1. Obtaining a closed *Ca*. Odinarchaeum LCB_4 genome.** (A) Summary methodology for the reassembly, refinement and closing of the *Ca*. Odinarchaeum LCB_4 genome. (B) Schematic of the assembly status before long-range PCR, indicating the presence of gaps and the agreement between two separate assemblies, which guided primer design. (C) Purified PCR products; lane 1: DNA ladder, 2: Positive control ca. 5 kb rRNA gene cluster; 3: Positive control ca. 10 kb ribosomal protein gene cluster; 4-5: first gap closing, at distances of ca. 5 and 5.5 kb; 6-8: second gap closing, at distances of ca. 4, 4.5 and 5 kb. (D) Genome map of *Ca*. Odinarchaeum LCB_4, including (from inside out): GC skew (line) and cumulative GC skew (histogram); GC content; Crick strand genes; Watson strand genes; Nanopore reads coverage capped at 1500X; Illumina read coverage (light: proper pairs, NM<3) capped at 50X; repeats; chromosome. (E) Comparison between previous assembly and new assembly for Huginnvirus, indicating circularity. Similarity lines represent two single BlastN hits with up to 1 mismatches. (F) Genomic patterns of the *Ca*. Odinarchaeum LCB_4 indicating a potential origin of replication at position 959350.

**Supplementary Figure S2. Predicted structure of selected proteins.** Comparisons between the structures of (A) DJR-MCPs (left: Huginnvirus: OLS18934.1; right: Sulfolobus turreted icosahedral virus 1: 3J31); (B) His1-like MCPs (left: Muninnvirus: OLS18630.1; right: His1 virus: YP_529533.1); (E) SIRV2-like MCPs (left: Jordarchaeia QZMA23B3: QZMA23B3_25900; right: Sulfolobus islandic*us* rod-shaped virus 2 (SIRV-2): 3J9X) and (F) transmembrane proteins (left: Muninnvirus: OLS18631.1; right: Ca. Odinarchaeum LCB_4 virus: OLS16720). All structures predicted with RoseTTAFold are color-coded according to their error estimate (Å). (C,D) Given the high error estimates for the predicted structures of His1-like MCPs (B), we append HHsearch results for OLS18630.1 (Muninnvirus) and OLS18934.1 (*Ca*. Odinarchaeote LCB_4 MAG), the latter of which shows a tandem duplication (Regions 1 and 2) of the His1-like MCP. H(h), α-helix; E(e), β-strand; C(c), coil.

**Supplementary Figure S3. Taxonomic placement of archaeal MAGs.** Phylogenomic tree obtained with FastTree including two archaeal MAGs (arrows) containing viral contigs within all current GTDB Archaea representatives (see methods). Branch colors represent Nitrososphaeria (red), Asgard archaea (light blue), and Jordarchaeia (purple). All placements are supported with branch support values of 1.0. Full tree can be found in data repository.

**Supplementary Figure S4. PolB phylogeny.** Midpoint-rooted full tree corresponding to Fig. 2A. Taxon labels shown in Fig 2A are colored with the same pattern, and their corresponding branches are colored red. Support values are transfer bootstrap expectation (left) and Felsenstein bootstrap proportions (right).

## REFERENCES

1 Zaremba-Niedzwiedzka, K. et al. Asgard archaea illuminate the origin of eukaryotic cellular complexity. Nature 541, 353–358, doi:10.1038/nature21031 (2017).

2 Williams, T. A., Cox, C. J., Foster, P. G., Szollosi, G. J. & Embley, T. M. Phylogenomics provides robust support for a two-domains tree of life. Nat Ecol Evol 4, 138–147, doi:10.1038/s41559-019-1040-x (2020).

3 Eme, L., Spang, A., Lombard, J., Stairs, C. W. & Ettema, T. J. G. Archaea and the origin of eukaryotes. Nat Rev Microbiol 15, 711–723, doi:10.1038/nrmicro.2017.133 (2017).

4 Eme, L. et al. Inference and reconstruction of the Skadiarchaeial ancestry of eukaryotes. Manuscript under revision (2021).

5 Farag, I. F., Zhao, R. & Biddle, J. F. “Sifarchaeota,” a Novel Asgard Phylum from Costa Rican Sediment Capable of Polysaccharide Degradation and Anaerobic Methylotrophy. Appl Environ Microbiol 87, doi:10.1128/AEM.02584-20 (2021).

6 Liu, Y. et al. Expanded diversity of Asgard archaea and their relationships with eukaryotes. Nature 593, 553–557 (2021).

7 Sun, J. E., P.N.; Gagen, E.J.; Woodcroft, B.J.; Hedlund, B.P.; Woyke, T.; Hugenholtz, P.; Rinke, C.. Recoding of stop codons expands the metabolic potential of two novel Asgardarchaeota lineages. ISME Communications 1, 30 (2021).

8 Spang, A. et al. Complex archaea that bridge the gap between prokaryotes and eukaryotes. Nature 521, 173–179, doi:10.1038/nature14447 (2015).

9 Yutin, N., Backstrom, D., Ettema, T. J. G., Krupovic, M. & Koonin, E. V. Vast diversity of prokaryotic virus genomes encoding double jelly-roll major capsid proteins uncovered by genomic and metagenomic sequence analysis. Virol J 15, 67, doi:10.1186/s12985-018-0974-y (2018).

10 Krupovic, M., Quemin, E. R., Bamford, D. H., Forterre, P. & Prangishvili, D. Unification of the globally distributed spindle-shaped viruses of the Archaea. J Virol 88, 2354–2358, doi:10.1128/JVI.02941-13 (2014).

11 Krupovic, M., Cvirkaite-Krupovic, V., Iranzo, J., Prangishvili, D. & Koonin, E. V. Viruses of archaea: Structural, functional, environmental and evolutionary genomics. Virus Res 244, 181–193, doi:10.1016/j.virusres.2017.11.025 (2018).

12 Wang, H. et al. Novel Sulfolobus Virus with an Exceptional Capsid Architecture. J Virol 92, doi:10.1128/JVI.01727-17 (2018).

13 Krupovic, M. et al. Adnaviria: a New Realm for Archaeal Filamentous Viruses with Linear A-Form Double-Stranded DNA Genomes. J Virol 95, e0067321, doi:10.1128/JVI.00673-21 (2021).

14 Ofir, G. et al. DISARM is a widespread bacterial defence system with broad anti-phage activities. Nat Microbiol 3, 90–98, doi:10.1038/s41564-017-0051-0 (2018).

15 Bernheim, A. et al. Prokaryotic viperins produce diverse antiviral molecules. Nature 589, 120–124, doi:10.1038/s41586-020-2762-2 (2021).

16 Doron, S. et al. Systematic discovery of antiphage defense systems in the microbial pangenome. Science 359, doi:10.1126/science.aar4120 (2018).

17 Baker, B. J. et al. Genomic inference of the metabolism of cosmopolitan subsurface Archaea, Hadesarchaea. Nat Microbiol 1, 16002, doi:10.1038/nmicrobiol.2016.2 (2016).

18 Li, H. Minimap2: pairwise alignment for nucleotide sequences. Bioinformatics 34, 3094–3100, doi:10.1093/bioinformatics/bty191 (2018).

19 Wick, R. R., Judd, L. M., Gorrie, C. L. & Holt, K. E. Unicycler: Resolving bacterial genome assemblies from short and long sequencing reads. PLoS Comput Biol 13, e1005595, doi:10.1371/journal.pcbi.1005595 (2017).

20 Li, D. et al. MEGAHIT v1.0: A fast and scalable metagenome assembler driven by advanced methodologies and community practices. Methods 102, 3–11, doi:10.1016/j.ymeth.2016.02.020 (2016).

21 Langmead, B. & Salzberg, S. L. Fast gapped-read alignment with Bowtie 2. Nat Methods 9, 357–359, doi:10.1038/nmeth.1923 (2012).

22 Biswas, A., Staals, R. H., Morales, S. E., Fineran, P. C. & Brown, C. M. CRISPRDetect: A flexible algorithm to define CRISPR arrays. BMC Genomics 17, 356, doi:10.1186/s12864-016-2627-0 (2016).

23 Padilha, V. A., Alkhnbashi, O. S., Shah, S. A., de Carvalho, A. & Backofen, R. CRISPRcasIdentifier: Machine learning for accurate identification and classification of CRISPR-Cas systems. Gigascience 9, doi:10.1093/gigascience/giaa062 (2020).

24 Galperin, M. Y., Makarova, K. S., Wolf, Y. I. & Koonin, E. V. Expanded microbial genome coverage and improved protein family annotation in the COG database. Nucleic Acids Res 43, D261–269, doi:10.1093/nar/gku1223 (2015).

25 Camacho, C. et al. BLAST+: architecture and applications. BMC Bioinformatics 10, 421, doi:10.1186/1471-2105-10-421 (2009).

26 Jones, P. et al. InterProScan 5: genome-scale protein function classification. Bioinformatics 30, 1236–1240, doi:10.1093/bioinformatics/btu031 (2014).

27 Steinegger, M. et al. HH-suite3 for fast remote homology detection and deep protein annotation. BMC Bioinformatics 20, 473, doi:10.1186/s12859-019-3019-7 (2019).

28 Mistry, J. et al. Pfam: The protein families database in 2021. Nucleic Acids Res 49, D412–D419, doi:10.1093/nar/gkaa913 (2021).

29 Burley, S. K. et al. RCSB Protein Data Bank: powerful new tools for exploring 3D structures of biological macromolecules for basic and applied research and education in fundamental biology, biomedicine, biotechnology, bioengineering and energy sciences. Nucleic Acids Res 49, D437–D451, doi:10.1093/nar/gkaa1038 (2021).

30 Chandonia, J. M., Fox, N. K. & Brenner, S. E. SCOPe: classification of large macromolecular structures in the structural classification of proteins-extended database. Nucleic Acids Res 47, D475–D481, doi:10.1093/nar/gky1134 (2019).

31 Lu, S. et al. CDD/SPARCLE: the conserved domain database in 2020. Nucleic Acids Res 48, D265–D268, doi:10.1093/nar/gkz991 (2020).

32 UniProt, C. UniProt: the universal protein knowledgebase in 2021. Nucleic Acids Res 49, D480–D489, doi:10.1093/nar/gkaa1100 (2021).

33 Guy, L., Kultima, J. R. & Andersson, S. G. genoPlotR: comparative gene and genome visualization in R. Bioinformatics 26, 2334–2335, doi:10.1093/bioinformatics/btq413 (2010).

34 Baek, M. et al. Accurate prediction of protein structures and interactions using a three-track neural network. Science 373, 871–876, doi:10.1126/science.abj8754 (2021).

35 Kim, J. G. et al. Spindle-shaped viruses infect marine ammonia-oxidizing thaumarchaea. Proc Natl Acad Sci U S A 116, 15645–15650, doi:10.1073/pnas.1905682116 (2019).

36 Schaffer, A. A. et al. Improving the accuracy of PSI-BLAST protein database searches with composition-based statistics and other refinements. Nucleic Acids Res 29, 2994–3005, doi:10.1093/nar/29.14.2994 (2001).

37 Fu, L., Niu, B., Zhu, Z., Wu, S. & Li, W. CD-HIT: accelerated for clustering the next-generation sequencing data. Bioinformatics 28, 3150–3152, doi:10.1093/bioinformatics/bts565 (2012).

38 Katoh, K. & Standley, D. M. MAFFT multiple sequence alignment software version 7: improvements in performance and usability. Mol Biol Evol 30, 772–780, doi:10.1093/molbev/mst010 (2013).

39 Capella-Gutierrez, S., Silla-Martinez, J. M. & Gabaldon, T. trimAl: a tool for automated alignment trimming in large-scale phylogenetic analyses. Bioinformatics 25, 1972–1973, doi:10.1093/bioinformatics/btp348 (2009).

40 Minh, B. Q. et al. IQ-TREE 2: New Models and Efficient Methods for Phylogenetic Inference in the Genomic Era. Mol Biol Evol 37, 1530–1534, doi:10.1093/molbev/msaa015 (2020).

41 Kalyaanamoorthy, S., Minh, B. Q., Wong, T. K. F., von Haeseler, A. & Jermiin, L. S. ModelFinder: fast model selection for accurate phylogenetic estimates. Nat Methods 14, 587–589, doi:10.1038/nmeth.4285 (2017).

42 Wang, H. C., Minh, B. Q., Susko, E. & Roger, A. J. Modeling Site Heterogeneity with Posterior Mean Site Frequency Profiles Accelerates Accurate Phylogenomic Estimation. Syst Biol 67, 216–235, doi:10.1093/sysbio/syx068 (2018).

43 Lemoine, F. et al. Renewing Felsenstein’s phylogenetic bootstrap in the era of big data. Nature 556, 452–456, doi:10.1038/s41586-018-0043-0 (2018).

44 Parks, D. H. et al. A complete domain-to-species taxonomy for Bacteria and Archaea. Nat Biotechnol 38, 1079–1086, doi:10.1038/s41587-020-0501-8 (2020).

45 Zhang, J. W. et al. Newly discovered Asgard archaea Hermodarchaeota potentially degrade alkanes and aromatics via alkyl/benzyl-succinate synthase and benzoyl-CoA pathway. ISME J 15, 1826–1843, doi:10.1038/s41396-020-00890-x (2021).

46 Lee, M. D. GToTree: a user-friendly workflow for phylogenomics. Bioinformatics 35, 4162–4164, doi:10.1093/bioinformatics/btz188 (2019).

