## Supplementary material for "A closed *Candidatus* Odinarchaeum genome exposes Asgard archaeal viruses": FigS2.pdf

A

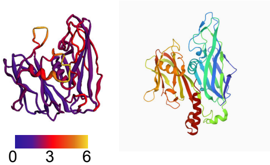


B

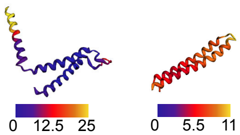


C

```
>Q25BH4 CAPSD_HIS1I Major capsid protein OS=His1 virus (isolate Australia/Victoria) OX=654912 GN=ORF21 PE=1 SV=1
Probab=99.33  E-value=1.2e-11  Score=85.37  Aligned_cols=63  Identities=27%  Similarity=0.452  Sum_probs=56.2
Template_Neff=3.400

Q ss_pred                cHHHHHHHHHHHHHHHHH----HHHHHHHHHHhccCceeeeCCCccchhhcCCHHHHHHHHHHHHHHHHHHHHHHhCcccch
Q Q_7599238            2 EIKYIVGLAISLIIIAV----IVPIGLAYLAGAANMSITVGNTTKTLSQWIDPTVLTLLTVLVPIMVVVGIVVAFVPKTSF  78 (99)
Q Consensus            2 ~i~~VI~iaV~L~Viai----ilPlgl~~la~a~da~itv~~~t~~l~t~vDpsVlTllTVLlPILaVIGIa~~Flp~~~~  78 (99)
                         +.+++|..++++++.++    ++|+++.+++   ++|.           |+||+++.|+.+ ++|+++++|++++|+|+...
T Consensus          3 ~l~~~V~~~VgllV~ailaA~L~Piai~~i-----vnT----------TgW~s~~~tiw~-lLpil~Vigv~l~fv~~a~~  69 (73)
T Q25BH4             3 EIDDGVEMAVGIFVVIILAANLLPTAFDQIF---NAST----------SSWNSDVTTLWE-LLPLLSVVGLLLMFVYWARK  69 (73)
T ss_pred                cHHHHHHHHHHHHHHHHHHHhHHHHHHHHH---cCCC----------CCCcHHHHHHH-HHHHHHHHHHHHHHHHHHHH
Confidence             46788999999986665    9999999999   7776          999999999999 99999999999999998753
```

D

```
Region 1.

>Q25BH4 CAPSD_HIS1I Major capsid protein OS=His1 virus (isolate Australia/Victoria) OX=654912 GN=ORF21 PE=1 SV=1
Probab=99.78  E-value=7.6e-19  Score=126.67  Aligned_cols=63  Identities=35%  Similarity=0.671  Sum_probs=54.3
Template_Neff=3.400

Q ss_pred                HHHHHHHHHHHHHHHHHHHHHHHHHHHHHccCCCCCcHHHHHHHHHHHHHHHHHHHHHHHHhccccCCC
Q Q_5042214          4 VNKAITIGVSLVILGAVLPTAIFTLIATDTSNWNSAVATIYSVVLPLLAVIGIVFWFMPRSKRG   67 (145)
Q Consensus          4 m~kIVgV~VGLLLAAiLlPiAI~~I~~vNTTGWDsTV~TIWtVLLPIL~VIGV~L~Fvk~s~~G   67 (145)
                       ++.++|++++.+++|+++|++++|++++|||||++++|+|+ +||+++++|++++|+|+.++.
T Consensus          8 V~~~VgllV~ailaA~L~Piai~~i~~vnTTgW~s~~~tiw~-lLpil~Vigv~l~fv~~a~~~   70 (73)
T Q25BH4           8 VEMAVGIFVVIILAANLLPTAFDQIFNASTSSWNSDVTTLWE-LLPLLSVVGLLLMFVYWARKA   70 (73)
T ss_pred                HHHHHHHHHHHHHHHHhhHHHHHHHHHHcCCCCCCcHHHHHHH-HHHHHHHHHHHHHHHHHHHHHc
Confidence             3444555555557777999999999999999999999999 99999999999999998753


Region 2.

>Q25BH4 CAPSD_HIS1I Major capsid protein OS=His1 virus (isolate Australia/Victoria) OX=654912 GN=ORF21 PE=1 SV=1
Probab=96.39  E-value=0.072  Score=40.28  Aligned_cols=62  Identities=23%  Similarity=0.456  Sum_probs=0.0  Template_Neff=3.400

Q ss_pred                HHHHHHHHHHH----HHHHHhhhcHHHHhcCCcceecccccceecCHHHHHHHHHhHHHHHHHHHHHHHHHHHHhHHH
Q Q_5042214         72 SRVIQIAVVLM----IVSALVPYGISQLASMPTATTVGTNTYTVNPAVLTMLKTVTPILAIIGIALFFLTVTASK  142 (145)
Q Consensus         72 ~~~~~~~~~~~----~~~~~~~~~~~~~~~~~~~~~~~~~~~~~~~~~~~~~~~~~~~~~~~~~~~~~~~~~~~  142 (145)
                       ++++..++.+.    +.+++.|-+++|+.        +.||-..|.++.|... +.|+++++|+++.|+....++
T Consensus          5 ~~~V~~~VgllV~ailaA~L~Piai~~i---------~vnTTgW~s~~~tiw~-lLpil~Vigv~l~fv~~a~~~   70 (73)
T Q25BH4           5 DDGVEMAVGIFVVIILAANLLPTAFDQIF--------NASTSSWNSDVTTLWE-LLPLLSVVGLLLMFVYWARKA   70 (73)
T ss_pred                HHHHHHHHHHHHHHHHHHhhHHHHHHHHHH--------cCCCCCCcHHHHHHH-HHHHHHHHHHHHHHHHHHHHHc
```

E

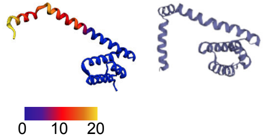


F

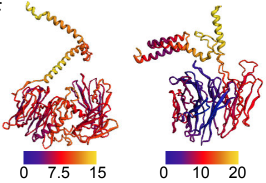
