## Supplementary figures and images for "A closed *Candidatus* Odinarchaeum genome exposes Asgard archaeal viruses"

### FigS1.pdf

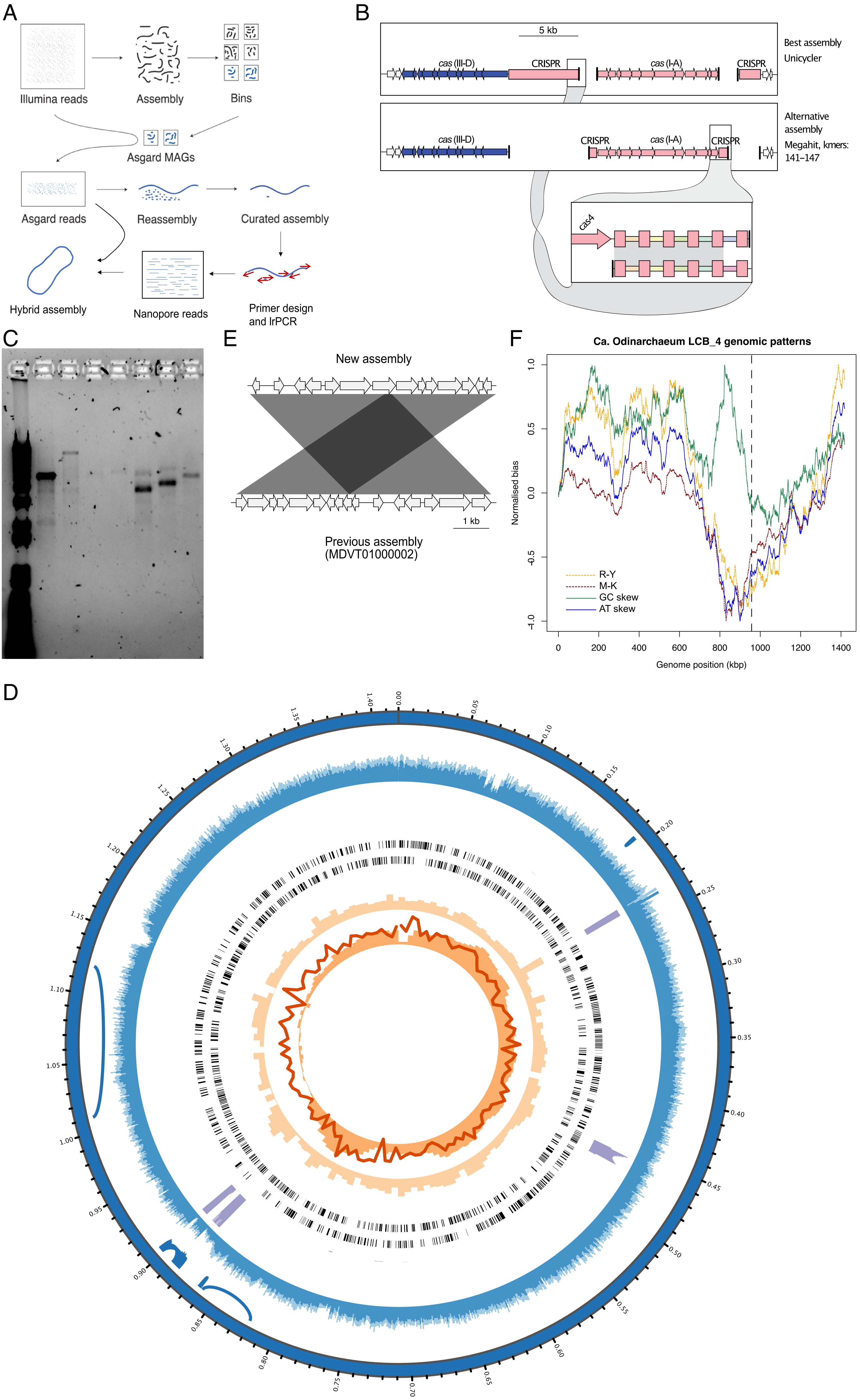

### FigS3.pdf

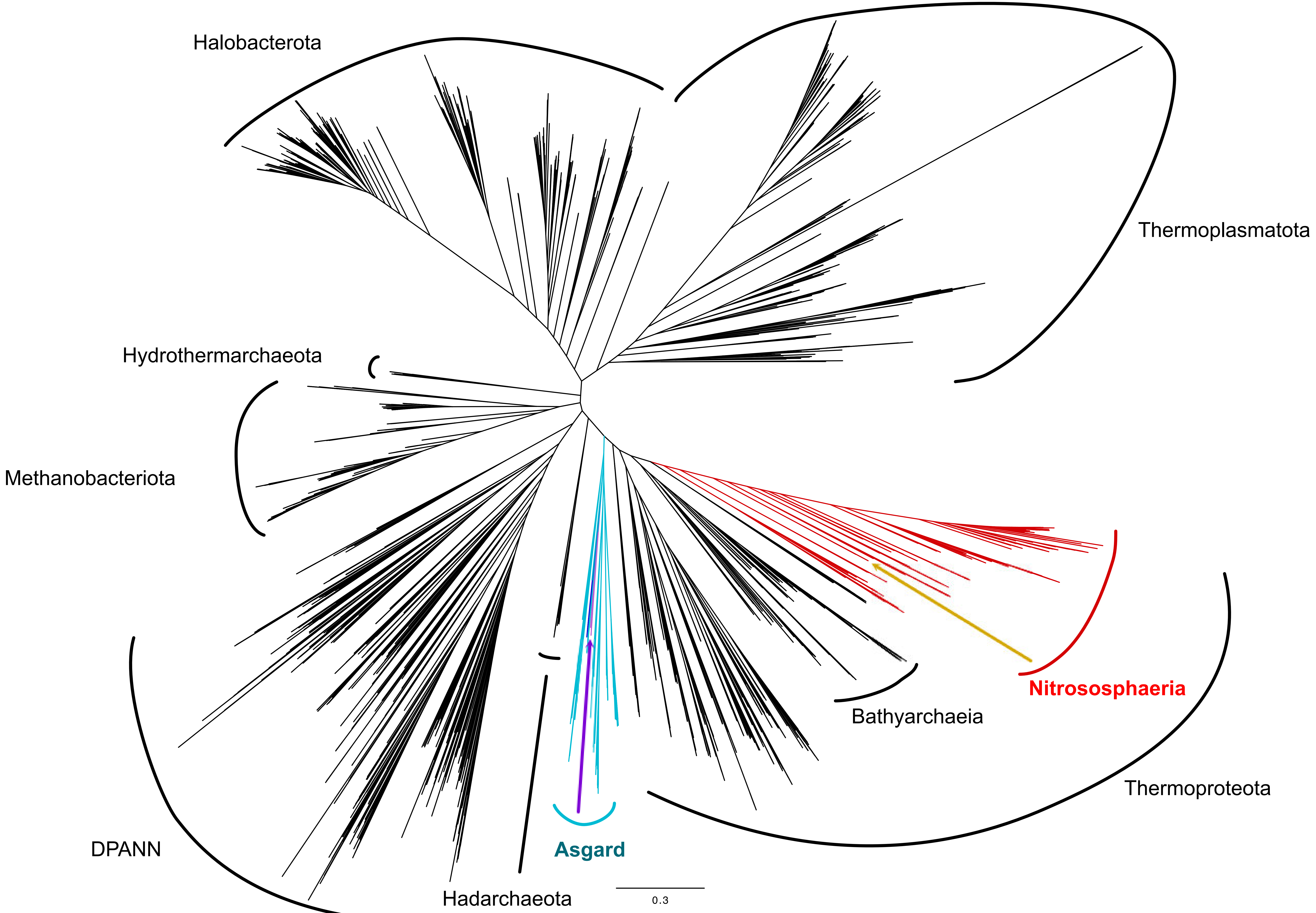

### FigS4.pdf

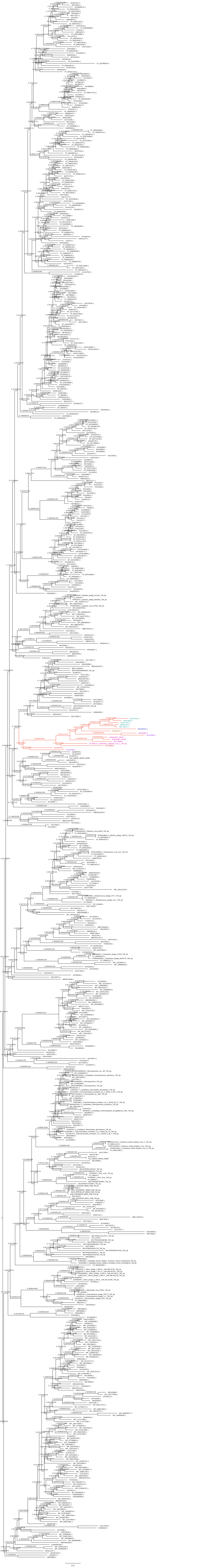
